# Uncovering Genes Essential in Domestication and Breeding of Sugar Beet

**DOI:** 10.1101/2024.12.02.626358

**Authors:** Amar Singh Dhiman, Nazgol Emrani, Eva Holtgrewe-Stukenbrock, Mark Varrelmann, Christian Jung

**Affiliations:** Plant Breeding Institute, Christian-Albrechts-University of Kiel, Olshausenstr. 40, D-24098 Kiel, Germany; Max Planck Institute for Evolutionary Biology, August-Thienemann-Str. 2 24306 Plön, Germany; Botanical Institute, Christian-Albrechts-University of Kiel, Am Botanischen Garten 1-9 24118 Kiel, Germany; Institute of Sugar Beet Research, Holtenser Landstr. 77, 37079 Göttingen

**Keywords:** *Beta vulgaris*, GWAS, population structure, mixed linear model, monogermity, bolting time, *Cercospora* leaf spot

## Abstract

The genus *Beta* encompasses economically important root crops such as sugar and table beet. A *Beta* diversity set including the wild relative *B. vulgaris* ssp. *maritima* was grown in the field, and a large phenotypic diversity was observed. The genomes of 290 accessions were sequenced, and more than 10 million high-quality SNPs were employed to study genetic diversity. A genome-wide association study was performed, and marker-trait associations were found for nine phenotypic traits. The candidate gene within the *M* locus controlling monogermity on chromosome 4 was previously unknown. The most significant association for monogermity was identified at the end of chromosome 4. Within this region, a non-synonymous mutation within the Zinc-Finger domain of the *WIP2* gene co-segregated with monogermity. This gene plays a regulatory role in *AGL8*/*FUL* in Arabidopsis. Intriguingly, commercial hybrids are in a heterozygous state at this position. Thus, the long-sought gene for monogermity was identified in this study. Red and yellow pigmentation due to betalain accumulation in shoots and roots is an important characteristic of table and leaf beets. The strongest associations were found upstream or downstream of two genes encoding Cytochrome P450 and anthocyanin MYB-like transcription factor proteins involved in betalain biosynthesis. Significant associations for *Cercospora* leaf spot resistance were identified on chromosomes 1, 2, 7, and 9. The associated regions harbor genes encoding proteins with leucine-rich repeats and nucleotide binding sites whose homologs are major constituents of plant-pathogen defense.

## 1. Introduction

Cultivated beets (*Beta vulgaris* ssp. *vulgaris* L.) are important crops belonging to the order Caryophyllales within the family Amaranthaceae. This group includes sugar, table, leaf, and fodder beets. It is widely accepted that all cultivated beet lineages originated from the wild progenitor *B*. *vulgaris* subsp. *maritima*, also known as ‘sea beet’.

Sugar beets are the second most important sugar-producing crop, contributing about 13% of the world’s sugar and the only source of sucrose in temperate regions^1^. They produce a thick, conical, fleshy taproot that accumulates sucrose up to 21% sucrose of its total fresh weight [1]. Table beets yield thickened roots and hypocotyls with intense red or yellow color, making them suitable for consumption as cooked vegetables. The variation in root colors is due to varying concentrations of the red-coloring pigment betacyanin and yellow-coloring pigment betaxanthin classified as betalains [2]. Owing to these compounds’ antioxidant properties, the demand for table beets is increasing as a food source and a natural red pigment utilized in the food industry [2–4].

Cultivated beets play integral roles across diverse sectors, including the global sugar industry, culinary arts, medicinal applications, and animal husbandry. The distinct characteristics and applications of each beet lineage underscore their economic, nutritional, and cultural significance on a global scale.

Leaf beets are consumed as a raw salad crop due to their succulent leaves. All above-ground vegetative parts, including leaves, petioles, and midribs, are used in salads. Leaf beets were among the earliest to undergo domestication from their wild ancestors. Importantly, they produce fangy roots resembling those of sea beets, suggesting deliberate selection for succulent leaves. The leaves exhibit various shapes, structures, and colors, including light green, dark green, yellow, pink, and red, making them an excellent choice for ornamental purposes [2, 4]. Fodder beets are utilized as a forage crop due to their thickened roots and hypocotyls. The roots consist of 80-85% water and contain a substantial amount of sucrose ranging from 4 to 10% [5]. Fodder beets typically yield broad and enlarged roots, with leaves and roots used as feed for cattle and other livestock.

Sea beets belong to the primary gene pool of cultivated beets due to relatively weak reproductive barriers, facilitating homologous recombination [6]. This genetic proximity has played a crucial role in successful breeding endeavors, allowing the incorporation of genetic diversity from sea beets to enrich the genetic makeup of cultivated beet germplasm. Sea beets are regarded as the most crucial wild relative, primarily because of their high genetic and phenotypic diversity against various biotic stresses caused by viruses, fungi, cyst nematodes, and insects, as well as resilience to abiotic stresses such as salt and drought [7–10].

The linkage between two coloration loci (*R* and *Y*) and the locus controlling early bolting (*B*) was one of the first linkage relationships ever identified in a plant species. The *R-Y-B* linkage group was later mapped on chromosome 2. The *R* and *Y* loci control betalain accumulation in beets. Molecular studies revealed that the underlying gene at the *R* locus (*CYP76AD1)* encodes a novel cytochrome P450, which is required for red betacyanin pigments [11, 12]. The *Y* locus contains the anthocyanin MYB-like protein gene *BvMYB1*, determining the presence of pigments, whether red or yellow, in the flesh of beet roots [11, 12].

Wild beet seeds are complex clusters of 2 to 11 fruits called seed balls. Consequently, each seed ball gives rise to more than one seedling. Thus, the seeds are called ‘multigerm’. This characteristic is undesirable in modern beet cultivation because it makes sugar beet cultivation highly labor-intensive due to the need to thin out the extra seedlings for optimal root growth. The discovery of monogermity in beets revolutionized sugar beet cultivation worldwide. The locus (*M*) controlling monogermity has been mapped at the end of chromosome 4, and the monogerm trait is recessively inherited (*m*) [4, 13]. Modern varieties are genetically multigerm (*Mm*) but phenotypically monogerm because the seed is harvested from a monogerm seed parent (*mm*) pollinated by a multigerm (*MM*) pollinator.

*Cercospora* leaf spot (CLS), caused by the fungus *Cercospora beticola* Sacc., stands as the most destructive foliar pathogen impacting beets globally [14]. This disease poses a significant threat to beet cultivation, especially in regions with warm and humid climates. The fungus produces a low-molecular-weight photo-activated toxin called cercosporin. Cercosporin generates singlet oxygen upon exposure to light, causing the near-simultaneous cell collapse within the sizable area. Consequently, this process leads to the progressive destruction of leaves, followed by the emergence of new leaves at the expense of the sucrose stored in the roots [15, 16]. Under high disease pressure, sucrose recovery may be compromised, potentially resulting in up to a 50% loss in recoverable sucrose [15, 17, 18]. Furthermore, increased impurities because of CLS contribute to additional economic losses by complicating sucrose recovery processes, leading to increased processing costs and a reduction in the yield of extractable sucrose [19]. An important source of quantitative resistance in cultivated beets originates from wild sea beets collected from Italy [8]. The broad-sense heritability was estimated to be between 60-70% [20, 21], and numerous resistance QTL have been localized [22, 23] [24, 25].

The number of whole-genome level studies to reveal marker-trait associations is relatively low compared to other crops. Consequently, our understanding of the genetic basis of various important agronomic traits, including resistance to biotic stress, is limited. A recent GWAS study examined 13 crucial agronomic traits in 977 sugar beet accessions using 170,750 SNPs. One hundred fifty-nine significant SNPs associated with root and shoot traits were identified, and candidate genes were predicted [26]. The importance of considering the epistatic interactions between different loci to understand the genetic determinants of complex traits using association mapping was investigated in a previous study of 460 elite sugar beet inbred lines which were genotyped with 290 SNP markers [27]. Another association study based on a population of 924 elite sugar beet lines genotyped with 677 SNP markers also confirmed that both main effect QTL and epistatic QTL define the genetic architecture of important agronomic and physiological traits in sugar beet [28]. A recent GWAS study involving 328 beet accessions mapped genomic loci associated with drought tolerance [29].

This study assembled a *Beta* diversity set representing this species’s genetic and phenotypic diversity [30]. More than 10 million SNP markers were used in a genome-wide association study depicting QTL for agronomically important traits. Within these QTLs, candidate genes were localized with a putative function in storage root formation, sucrose storage, pathogen resistance, and seed morphology. These genes are proposed to be key genes for root crop domestication and sucrose storage.

## 2. Materials and Methods

### 2.1. Plant materials and growth conditions

We have assembled a *Beta* mini-core collection (BMC) representing the phenotypical and geographical diversity of *B. vulgaris* [30]. It consists of 290 different sea beet and cultivated beet accessions, displaying a high degree of phenotypic homogeneity. The BMC was grown in 2021 (April 21 to September 29) and 2022 (April 20 to September 22) in the field near Göttingen, Germany. Seeds were hand-sown as single-row plots, aiming for seven plants per row in a randomized complete block design with three replication blocks. Each replication block contained 290 row plots, each representing a single accession. Subsequently, each replication block was divided into two sub-blocks, each measuring 13.5m × 8m. The distances between rows and between plants were set to 45 cm and 20 cm, respectively. In total, 876 row plots were sown across all three replication blocks.

Standard agronomic practices, including fertilizer application and weed control, were implemented. After sowing, 75 kg/ha nitrogen fertilizer was applied across the field. In the first week of July, a foliar application of 3 l/ha of Lebosol-Bor was carried out. Weeds were controlled using standard herbicides, with three applications: 1.5 l/ha Goltix Titan, 2 l/ha Betasana, and 0.5 l/ha Oblix were applied at two-week intervals from the second week of May to the second week of June.

### 2.2. Phenotyping and data analysis

Visual phenotyping was conducted per plant (Table 1, Figure 1 and Supplementary Figure 1). Leaf color and leaf hairiness were recorded at BBCH50 within 60 days after sowing. Leaf color variations include light green, green, a mixture of green leaf blades with red veins, marked as ‘mix,’ and entirely red with both leaf blade and veins in red color (Figure 1 and Supplementary Figure 1). The core collection exhibited a considerable variation in leaf hairiness. The bolting date was recorded for each accession as the day of visible stem elongation. The row plots were phenotyped every two weeks from the onset of bolting until the last week of the cultivation period in the field trial 20 weeks after sowing. Non-bolting plants were classified as biennials. We performed visual phenotyping for hypocotyl color, ranging from green, pink, red, and yellow. Accessions exhibiting heterogeneity for hypocotyl color were excluded from the GWAS.

**Figure 1:**
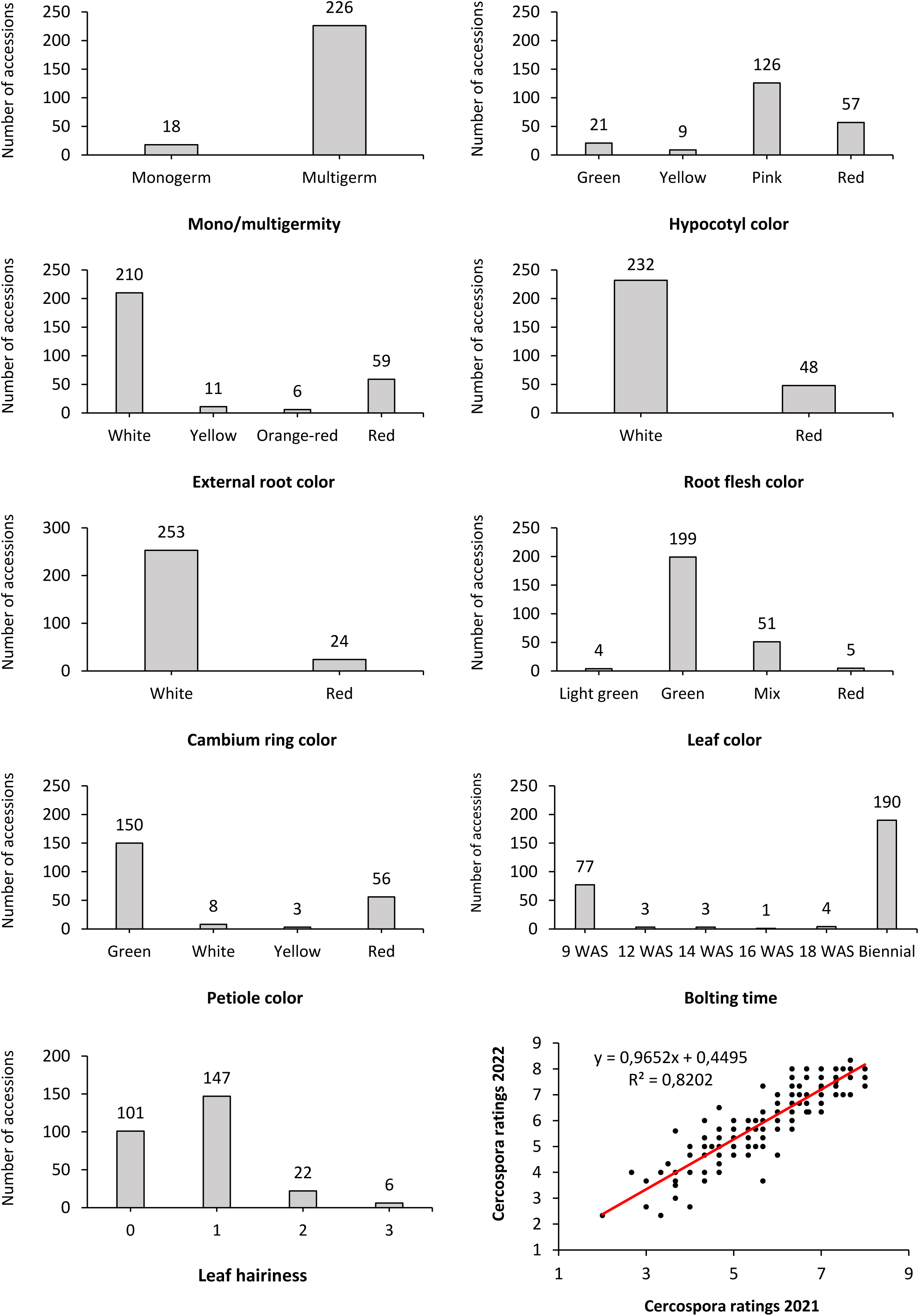
Frequency distribution plots illustrating phenotypic variation across the Beta mini-core collection. Plants were grown in the field in two consecutive years. Within each frequency distribution plot, the number of accessions belonging to each representative class is written on top of each vertical bar. The correlation of the Cercospora disease index, ranging from 1-9, is shown between 2021 and 2022 across 215 accessions.

**Table 1:**
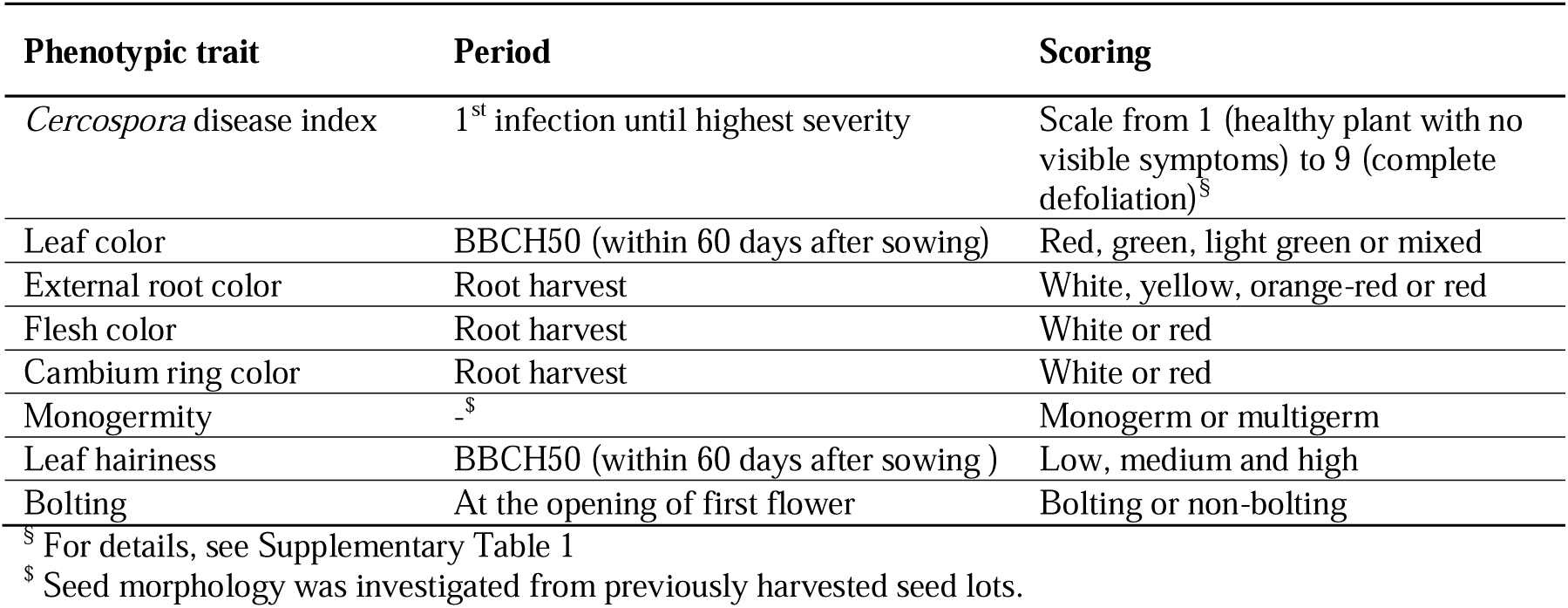
Phenotypic traits evaluated in this study.

Root, flesh, and cambium ring color were visually phenotyped at harvest. The root color ranged from white, yellow, orange-red, and red. The flesh color and cambium ring color of beetroots were assessed after cross-sectioning, and categorical white or red scores were assigned.

We characterized the seed type (multigerm or monogerm) by visual scoring the seeds used for the field trial. Monogerm seeds contain one embryo, while multigerm seeds contain more than one embryo. Visual phenotyping was applied to the seeds from each parent seed bag. Accessions with strictly monogram seeds and multigerm seeds were categorized as monogerm and multigerm, respectively. Only the phenotypic data from the accessions with a uniform seed type were used for GWAS. Sugar beet hybrid varieties were considered multigermic because they are produced by pollinating a monogerm seed parent with a multigerm pollinator.

### 2.3. *Cercospora* leaf spot infection screening

The field was artificially inoculated with dried CLS-infested leaf material collected from Hankensbüttel, Lower Saxony, Germany, to ensure uniform CLS infections. In brief, the collected leaf material was air-dried in an open hall. The following year, before the field inoculations, the dried leaves were hand-crushed to remove large chunks of leaf petioles. The crushed leaf material was sieved to remove the large chunks of leaf petioles. The ground leaf material was mixed with wheat semolina in a 1:5 w/w ratio and evenly dispersed across the field at a rate of 24g/m^2^ before sowing. Approximately 25 kg of the resulting mixture was applied to the field two days before sowing.

The first CLS screening was performed 12 weeks after sowing upon the first visible spots in the field. Intensive *Cercospora* screening was conducted every two weeks to examine the progression of disease severity over time. Six screenings were performed from the 12^th^ week, when the first spots became visible, until the 20^th^ week after sowing. The *Cercospora* disease index, measured on a scale of 1-9, was used to assess disease severity within each accession. A disease index rating of 1 indicates no visible symptoms, while 9 signifies complete defoliation. An increasing disease index represents higher *Cercospora* infection than preceding values (Supplementary Table 1, Figure 1).

The average disease index from three plots was used to calculate Best Linear Unbiased Estimators (BLUEs) with residuals approximately following a normal distribution. The statistical software R [31] was used for data analysis. The data evaluation began by defining an appropriate statistical mixed model [32, 33]. The model included the fixed factor ‘Genotype’. The ‘Year’ and the ‘Block,’ nested in ‘Year,’ were treated as random factors. Based on graphical residual analysis, residuals were assumed to follow an approximately normal distribution and to be heteroscedastic.

### 2.4. Genome-wide association study

The genotyping data from the *Beta* mini core collection [30] were used for the GWAS. In brief, paired-end (PE) short-reads were generated from an individual plant representing the whole accession using the Illumina NovaSeq 6000 sequencing platform. Raw reads were trimmed and filtered for bar-code adapters using Trimmomatic-v0.39 [34]. Standard pipelines were used for read alignment using BWA-MEM, sorting and indexing the intermediate files using SAMtools, and finally, GATK best practices were used to call and genotype the variants from 290 accessions onto a long-read genome assembly ‘EL10.2_2’ of a sugar beet inbred line EL10.

High-confidence SNPs with a minor allele frequency of 0.05 or greater were used for population structure, kinship, and GWAS analyses. A mixed-linear model (MLM) incorporating population structure and genetic relationships was employed to mitigate spurious associations and minimize false positives. The population structure (P matrix) within the *Beta* mini-core collection was estimated using the SNPrelate software [35]. The genetic relatedness among accessions, represented by a kinship matrix (K), was calculated using the Balding-Nicholas model in the efficient mixed-model association expedited (EMMAX) software [36]. The mixed linear model for the entire population, which included the K and P matrices, was employed in the EMMAX program for GWAS analyses. To determine genome-wide empirical significance thresholds, we applied Bonferroni correction with nominal levels of αL=L0.05 and αL=L1. This corresponded to raw *P* values of 4.82 × 10−9 at α = 0.05 and 9.65 × 10−8 at αL=L1, or −log_10_(*P*) values of 8.31 and 7.02, respectively. SNP *p*-values were plotted on Manhattan plots and quantile-quantile (Q-Q) plots using the ‘qqman’ package in R [37].

Trait-associated regions were defined by extending 100 kb upstream and downstream of the identified SNP markers. Overlapping regions were concatenated using bedtools (v2.26.0). To visualize highly associated variants within genic regions, we utilized the Integrative Genomics Viewer (IGV) v2.16.1[38] and the CLC workbench 7 (QIAGEN® Aarhus A/S, Aarhus C, Denmark). Furthermore, the domains within associated genes were predicted and plotted using the InterPro software tool [39].

## 3. Results

### 3.1. Phenotypic analysis of a Beta *mini*-core collection under field conditions

Initially, a diversity set of 385 *Beta* accessions was grown in row plots across three replication blocks in Kiel, Germany, in 2020. Those accessions displaying high phenotypic heterogeneity were discarded, resulting in 290 highly homogeneous accessions forming the *Beta* mini-core collection (BMC). It includes 91 sea beet (*B*. *vulgaris* ssp. *maritima*), 81 sugar beet, 59 table beet, 30 fodder beet, and 29 leaf beet accessions from 40 countries worldwide, effectively capturing extensive genetic diversity within the genus *B*. *vulgaris*. Within the BMS, cultivated beets display a spectrum of leaf colors, including light green, green, and red, as well as root colors spanning white, red, and yellow. The BMC integrates germplasm resources from various breeding programs, including those of the USDA, Italian beet germplasm resources, breeding companies, and genebank collections. The Western U.S. USDA breeding programs primarily focus on resistance to multiple biotic stresses like curly top, rhizomania, yellowing viruses, and mildew. The collection also showcases diversity in terms of monogerm and multigerm, as well as annual and biennial accessions.

The BMC displayed a huge phenotypic diversity when grown in the field for two consecutive years (Figure 1). Highly heritable characters were assessed. Out of 290 accessions, 226 accessions (77.93%) were homogenous for multigerm seeds, 46 accessions (15.86%) were segregating for mono- and multigerm seeds and only 18 accessions (6.20%) exhibited homogeneity for monogerm seeds. The BMC could be grouped into four classes, based on hypocotyl and external root color. In contrast, only two phenotypes were observed for root flesh and cambium ring color (white, red) (Supplementary Figure 1). All sugar beet accessions displayed green leaves and petioles, while most table beets had red leaf blades with red veins and petioles.

Bolting time was assessed until the end of the field trial. Of 290 accessions, 190 (65.52%) were biennial, and 88 (30.34%) were annuals.The remaining accessions were segregating for bolting. Bolting time varied among the annual accessions, ranging from 9 to 18 weeks after sowing (WAS) (Figure 1). As expected, most sugar, table, and fodder beet accessions were biennial.

This study focused on *Cercospora* Leaf Spot (CLS) infection. Homogeneous infections were observed across the replication blocks due to artificial inoculation using *Cercospora*-infected leaves. The first symptoms in 2021 and 2022 occurred 12 and 13 weeks after sowing. Seventy-five accessions were excluded from further CLS screenings due to their very early bolting, resulting in a smaller foliar size (Figure 1). The remaining accessions displayed CLS scores between 1 and 8 (Supplementary Figure 2). Table and fodder beet accessions exhibited higher disease index values in both years. The linear regression of *Cercospora* disease indexes across two years, involving 215 accessions, exhibited a strong Pearson correlation of 0.82 (Figure 1). Consequently, the averages derived from each plot in both years were employed to calculate the Best Linear Unbiased Estimates (BLUEs), used as phenotype values for the GWAS analysis. Heritability calculations from the replication blocks in both years resulted in a broad-sense heritability (h²) of 0.89 across the entire dataset. All phenotypic data are presented in Supplementary Table 2.

### 3.2. Genetic diversity and Kinship relationship among accessions of the Beta diversity panel

Approximately 11.7 million variants, including SNPs (10,365,662) and InDels (1,303,892), were genotyped (Figure 2A), with an average of 20.5 variants/kb. Notably, regions near the chromosome ends exhibited a higher density of variants. Regions devoid of variants correspond to ‘Ns’ stretches in the ‘EL10.2_2’ genome assembly. Interestingly, certain regions across nine beet chromosomes showed fewer variants across the 290 wild sea beet and cultivated beet accessions. Given a substantial density of SNPs/kb on all nine chromosomes, averaging 18.22 SNPs/kb, only High-Confidence SNPs (HC-SNPs), excluding InDels, were used for downstream analyses. This comprehensive genotypic characterization lays the foundation for exploring genetic associations within the wild and cultivated beet collection.

**Figure 2:**
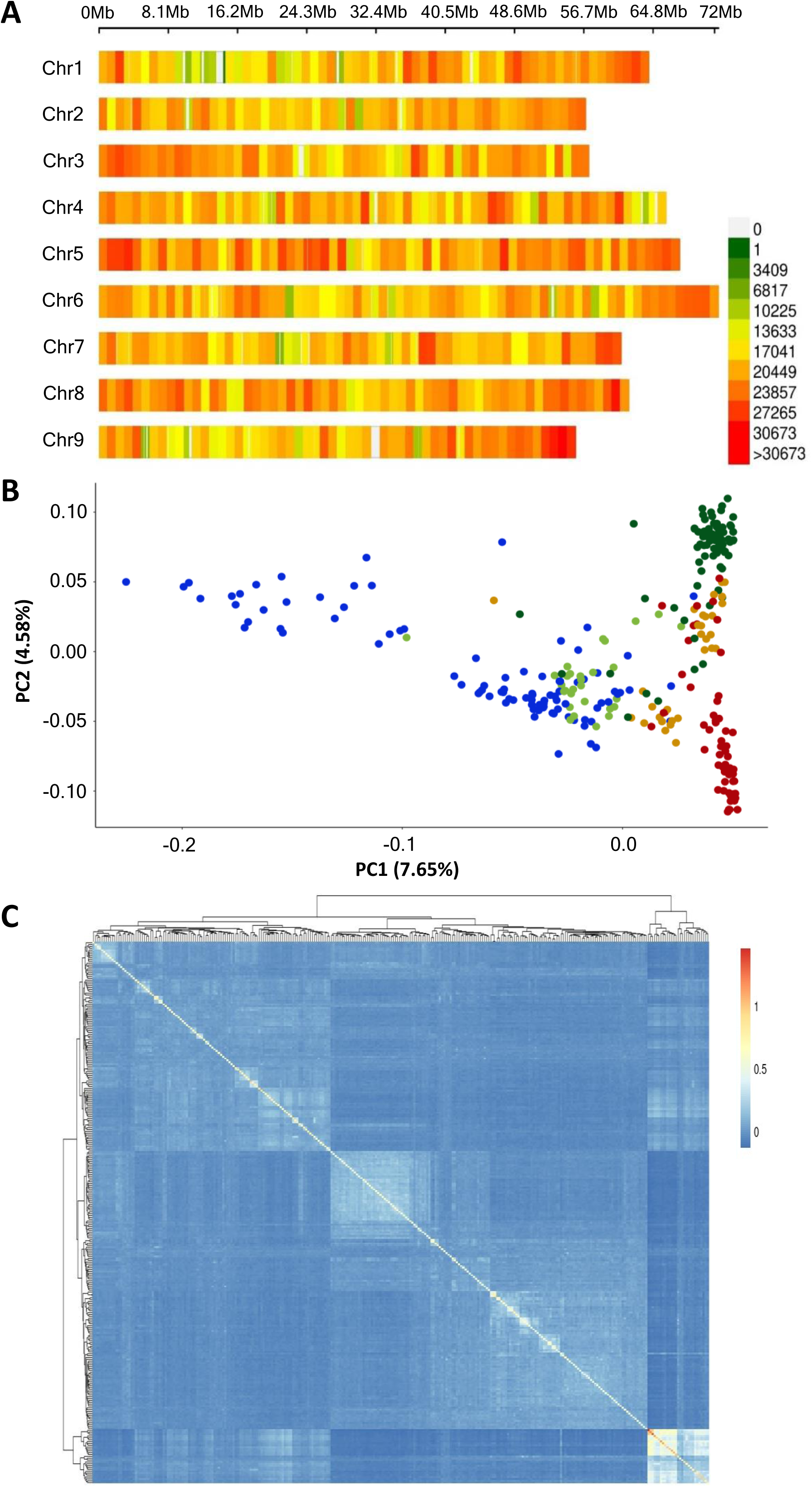
Distribution of variants, population structure, and genetic relationships among 290 accessions in the Beta mini-core collection. (A) The plot illustrates the variant distribution across all nine ‘EL10’ beet chromosomes depicted in a density plot. The color shows the variant density in a 1 Mb window. CMplot package in R was used for plotting, with the top X-axis showing chromosome length in Mb. (B) Principal component analysis (PCA) of the 290 beet accessions, with PC1 and PC2 accounting for 7.65% and 4.58% of the total variation, respectively. Colors indicate different genetic clusters representing wild sea beet (blue), sugar beet (dark green), table beet (red), fodder beet (orange), and leaf beet (light green) accessions. (C) Kinship matrix of the 290 accessions based on the Balding-Nicolas model in EMMAX, plotted using the pheatmap function in R. The dendrogram depicts phylogenetic relationships, while color in the kinship matrix represents genetic relatedness between accessions in a heat map.

A Principal Component Analysis (PCA) was performed using HC-SNPs to calculate P-matrices for GWAS analyses and gain insights into population structure (Figure 2B). PC1, explaining 7.65% of the variance, differentiated sea beet accessions from the cultivated beet accessions. Furthermore, PC2, accounting for 4.58% of the variance, distinguished sugar beet from table beet accessions. The relatively lower variation explained by PC1 and PC2 indicates that the panel does not represent a highly structured population. The resulting P-matrix, derived from the first five principal components (PCs), was a cofactor in subsequent GWAS analyses.

Genetic relatedness between accessions was estimated through marker-based kinship using the Balding-Nicholas model EMMAX. Clustering analysis revealed the identification of two sub-clusters. Notably, sea beet accessions originating from the Atlantic region formed a distinct cluster separate from the other accessions. Most accessions exhibited lower kinship fg-values and, typically, less than 0.5. Sub-structuring based on different crop types was also evident (Figure **2**C). A kinship matrix (K-matrix) was incorporated into the GWAS analyses to mitigate the risk of false positives resulting from genetic relatedness.

### 3.3. Genome regions harboring candidate genes for agronomically important traits

We performed genome-wide association studies for ten agronomically important traits (Table 1) using phenotypic data from 290 accessions (Supplementary Table 2) and 10,365,662 SNP markers. We identified 2,002 highly significant marker-trait associations (MTAs) for all evaluated traits (Supplementary Table 3).

First, we used hypocotyl, leaf, and root coloration as a proof-of-principle experiment because the respective genes controlling color formation in beet had been cloned [11, 12]. The *R* locus harbors the *CYP76AD1* gene, spanning from position 49,286,874 to 49,291,710, whereas the *Y* locus contains the *BvMYB1* gene, spanning from position 51,654,941-51,658,812 encodes for a transcription factor for both *CYP76AD1* and *DODA1*. Both genes are located on chromosome 2 at a distance of ∼2.36 Mb. The strongest association for hypocotyl color was identified on chromosome 2 (Figure 3, Supplementary Table 4). While comparing plants with green and yellow hypocotyls, the significant markers co-localized exclusively with the *Y* locus. The significantly associated region spanned from position 51,508,888 to 51,602,121 on chromosome 2, ∼50 kb upstream of the *BvMYB1* gene (Supplementary Table 5). Then, we compared accessions with green and red hypocotyl. The significant markers co-localized exclusively with the *R* locus on chromosome 2 (Figure 3, Supplementary Table 6). The significantly associated region spanned from position 49,248,516 to 49,503,722. Interestingly, none of the associated SNPs were present within the ORF of the *CYP76AD1* gene (Supplementary Table 6).

**Figure 3:**
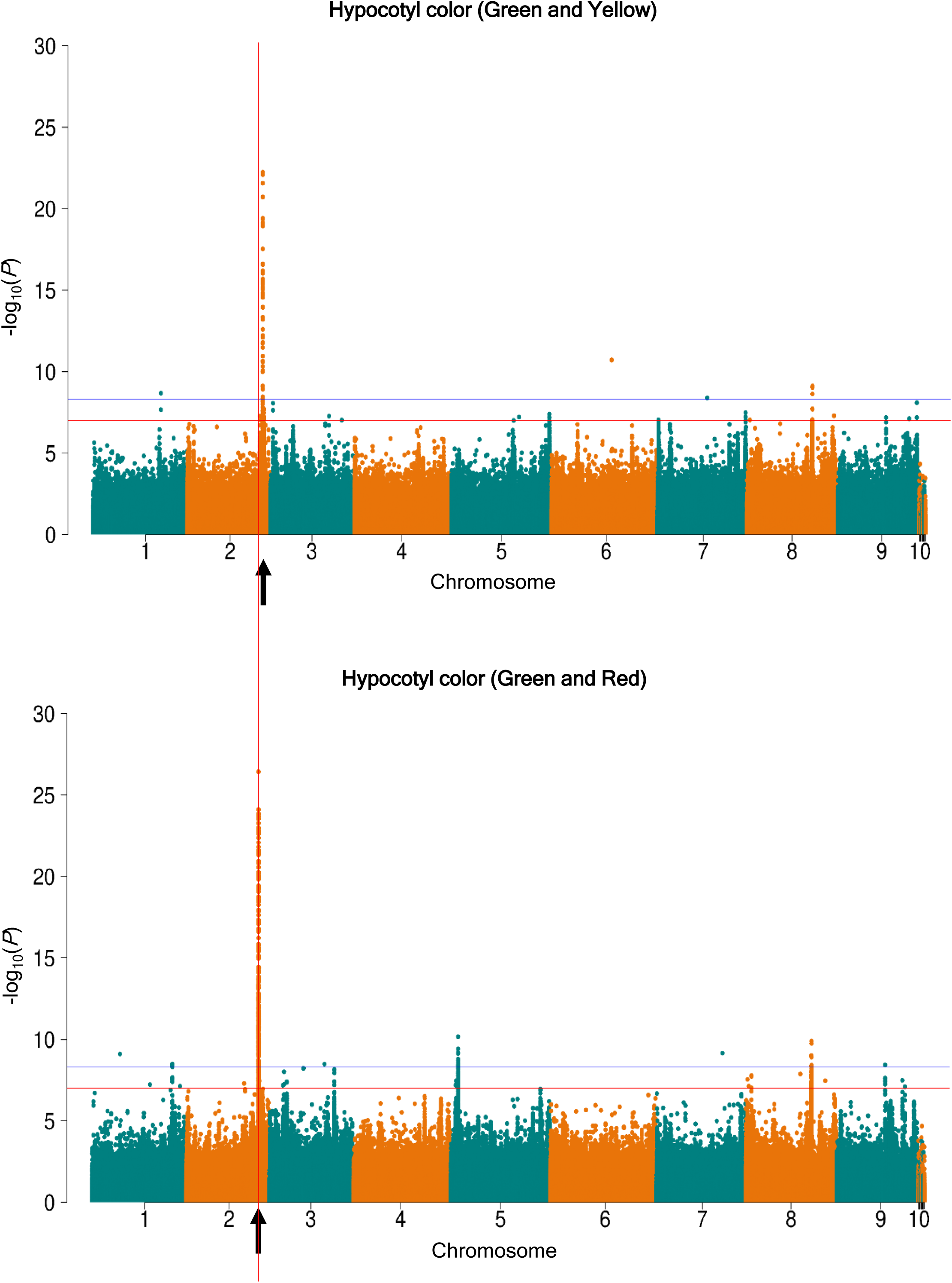
Manhattan plots to compare hypocotyl color using green and yellow and green and red as extremes. The GWAS for green and yellow revealed the most significant associations at the end of chromosome 2, co-localizing with the Y locus. However, when comparing green and red, the most significant associations were observed, co-localizing with the well-known R locus. In each Manhattan plot, the X-axis represents the nine main ‘EL10’ chromosomes and unplaced scaffolds labeled as 10, while the Y-axis illustrates the –log10(p) values. A vertical box was drawn to highlight the association for the R locus, which does not coincide with the Y locus. Two horizontal lines represent the Bonferroni thresholds of 7.02 (α=1) and 8.31 (α=0.05). The Manhattan plots were generated using the ‘qqman’ package in R. The arrows highlight the positions of the Y and R locus for green/yellow hypocotyl color and green/red hypocotyl color, respectively.

In the next step, we searched for marker-trait associations for red flesh color, external root color, leaf color, cambium ring color and petiole color (Supplementary Table 7-Supplementary Table **11**), which co-segregate with both *R* and *Y* loci [11, 12], suggesting the importance of both loci in controlling beet root pigmentation (Supplementary Figure 3 and Supplementary Figure 4). The highest number of associations was identified on chromosome 2 (Supplementary Table 4).

Specifically, 12 SNPs upstream and four SNPs downstream of the Cytochrome P450 gene ‘*CYP76AD1’* were significantly associated with the red root flesh color (Supplementary Table 7). However, there were no SNP associations within the *CYP76AD1’* ORF. Expectedly, we identified five SNPs within a gene ‘*DODA1’* encoding for 4, 5-DOPA-extradiol-dioxygenase, known to control the red pigmentation in beets [11, 40]. This gene is located ∼44 kb downstream of *CYP76AD1*. Additionally, 20 SNPs around the underlying genes, particularly *BvMYB1* within the *Y* locus, were significantly associated with red color (Supplementary Table 7).

Then, we compared accessions that differed in their external root color. Among the 171 MTAs, 150 were on chromosome 2 (Supplementary Table 4). The most significant MTAs were found near the *CYP76AD1, DODA1,* and *BvMYB1* genes, known for their role in determining the root color. Specifically, we identified 19 SNPs within the *CYP76AD1* ORF, which were reportedly required to produce red betacyanin pigments in beets [11]. Interestingly, the SNP associations around the *CYP76AD1* gene within the *R* locus had lower *P*-values than the SNP associations within the ORF of the gene (Supplementary Table 3 and Supplementary Table **8**).

Next, we searched for SNPs associated with monogermity. The locus had been located on chromosome 4 [13]. The most significant SNP at 64,590,348 bp is located within the coding region of the *BvWIP2* gene, which encodes a Zinc-finger protein (Figure 4, Supplementary Table 3 and Supplementary Table **12**). A missense mutation within the DNA-binding domain of the BvWIP2 protein co-segregates with mono- and multigermity in beets. Accessions with monogerm seeds carry the ‘*A*’ nucleotide, while those with multigerm seeds carry the*‘T’* nucleotide at this position (Supplementary Figure 5A). In sugar beet hybrid varieties, the SNP is present in the heterozygous state ‘AT,’ as expected. This suggests that *BvWIP2* is the long-sought gene responsible for monogermity in beets. BvWIP2 is 454 amino acids long, containing a classic C2H2 Zinc-finger domain.

**Figure 4:**
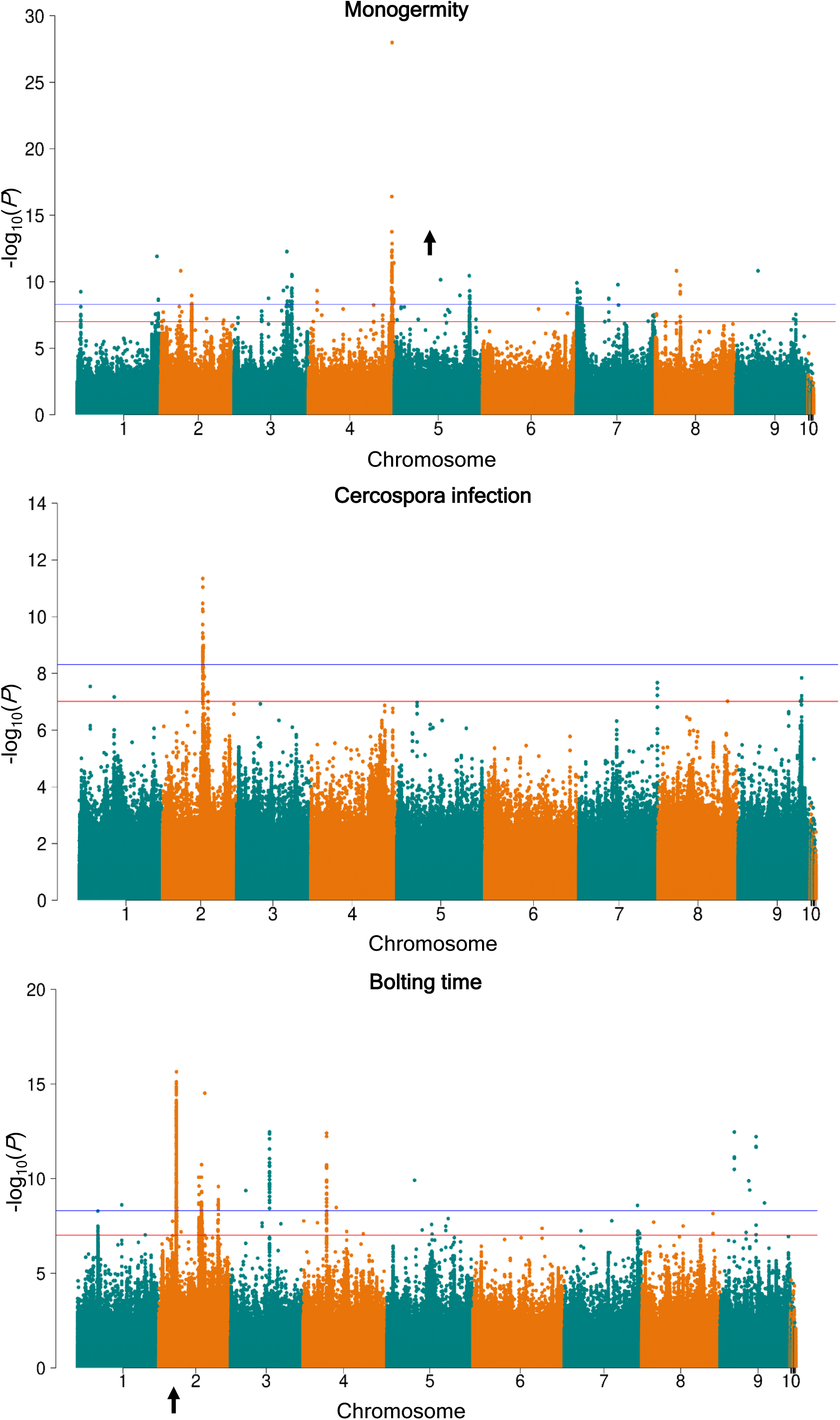
Marker-trait associations for monogermity, Cercospora resistance, and bolting time. A genome-wide association study was performed with phenotypic data from the Beta core collection. The most significant marker-trait associations are shown. In each Manhattan plot, the X-axis represents the nine main ‘EL10’ chromosomes and unplaced scaffolds labeled as 10, while the Y-axis illustrates the –log_10_(*p*) values. Two horizontal lines represent the Bonferroni thresholds of 7.02 (α=1) and 8.31 (α=0.05). The Manhattan plots were generated using the ‘qqman’ package in R. The arrows highlight the positions of the *M* and *B* locus for monogermity and bolting time, respectively.

Then, we investigated the association of the BvWIP2 protein in different biological, molecular, and cellular processes. According to InterPro [39] and PANTHER GO terms, the BvWIP2 protein is associated with the molecular function category ‘DNA-binding transcription factor activity (GO:0003700)’. Furthermore, the cellular component category shows associations ‘within the nucleus (GO:0005634)’. Importantly, in the biological process category, it is associated with the ‘regulation of gene expression (GO:0010468)’ and ‘anatomical structure development (GO:0048856)’ (Supplementary Figure 6). In Arabidopsis, *WIP2* plays a role in replum development by activating the homeobox protein KNAT1 [41], and its role in the regulation of *AGL8/FUL* has been speculated [42].

Finally, we identified MTAs for bolting time control. Most associations were found on chromosome 2 (Supplementary Table 4 and Supplementary Table **13**). We found 13 SNPs upstream of *BTC1* controlling the annual life cycle in *Beta* species [43]. However, the most significant region spanned from position 13,730,603 to 14,152,924, encompassing 373 SNPs significantly associated with bolting time control (Figure 4, Supplementary Table 3). Within this region, we identified a gene with high homology to an Arabidopsis gene, which is predicted to be involved in circadian rhythm regulation (ID ‘AT3G01060’) (Supplementary Table 13).

### 3.4. Genome regions and candidate genes associated with Cercospora leaf spot resistance

Highly significant marker-trait associations (MTAs) for CLS were identified on chromosomes 1, 2, 7, 8, and 9 (Figure 4, Supplementary Table 3, Supplementary Table **4** and Supplementary Table **14**). A previous QTL study mapped the most significant QTL for CLS tolerance to chromosome 2 [22]. Consistent with these findings, our study identified the strongest association on chromosome 2, from position 30,827,268 to 31,330,290. A gene putatively encoding an actin-depolymerizing factor 4 (ADF4) was identified within this region. The actin cytoskeleton is a vital component in plants, influencing the dynamic assembly of actin filaments. *ADF* genes primarily regulate the severing and depolymerization of actin filaments. ADFs are crucial in modulating the host’s cytoskeletal architecture and defense signaling during pathogen infection [44, 45].

Within the associated regions on chromosome 7, a cytochrome P450 gene (*CYP94C1*) was identified. In Arabidopsis, the *CYP94C1* homolog plays a role in the wound-induced accumulation of hydroxylated and di-carboxylated jasmonoyl-L-isoleucine (JA-Ile), crucial for oxidative turnover of the JA-Ile hormone [46].

In correspondence with previous studies, we identified another putative resistance locus on chromosome 9 [22, 23]. Within the associated region, we found a cluster of genes putatively belonging to the family of Leucine-Rich Repeat (LRR) protein kinase genes. LRR protein kinases are crucial in host-specific and non-host-specific defense response, wounding response, and symbiosis [47]. The LRR-containing domain in many proteins is evolutionarily conserved and associated with the plant’s innate immunity. LRR protein kinase genes serve as the first line of defense, initiating the innate immune response upon sensing pathogen-associated molecular patterns [48]. In summary, the vast majority of marker-CLS associations were on chromosome 2 (93), followed by chromosome 9 (5), chromosome 7 (4), chromosome 1 (2), and chromosome 8 (1). None of the associated SNPs were located within the ORF of the genes.

## 4. Discussion

Our recently established *Beta* mini-core collection displayed high phenotypic variation for agronomically relevant traits, including monogermity, Cercospora leaf spot (CLS) resistance, red pigmentation, trichome density (data not shown), and bolting control. The genotypic and phenotypic data were used to map genomic loci associated with the evaluated traits. We found significant marker-trait associations (MTA) for red pigmentation, CLS resistance, bolting control on chromosome 2, and monogermity on chromosome 4. This way, the long-sought candidate gene controlling monogermity has been identified. The deviation of *p*-values from the expected distribution in Q-Q plots for all the traits showed the number of confirmed associations deviating from the null hypothesis of no association (Supplementary Figure 7 and Supplementary Figure 8).

Our genome-wide study sheds light on genetic variation associated with red pigments in beets. The genes controlling red pigment production in beets had been cloned [11, 12]. The *R* locus harbors the gene *CYP76AD1*, which encodes a novel cytochrome P450 protein required to produce red betacyanin pigments in beets. When *CYP76AD1* is mutated, only yellow pigments are produced. The *Y* locus harbors the gene *BvMYB1*, which encodes an anthocyanin MYB-like transcription factor protein that regulates the betalain pathway by controlling the expression of both *CYP76AD1* and *DODA1* in beets. Both genes are located on chromosome 2 at a distance of ∼2.36 Mb. Our GWAS analyses revealed the strongest associations with red pigmentation in roots, leaves, and petioles in the non-translated regions up- and downstream of the *CYP76AD1* and *BvMYB1* genes. The significance values of MTAs within the genes were lower than MTAs around the genes, indicating that regulatory regions of these genes might play a crucial role in the production and regulation of red pigments in beets, as opposed to variations within the ORFs. It has also been shown that a 4, 5-DOPA-extradiol-dioxygenase (DODA1) plays a vital role in the betalain pathway [11, 40]. Interestingly, we identified five MTAs within a *DODA1* gene located within the associated region.

Species of the genus *Beta* display a broad variation in phenological development. Because the yield and quality of bolting and flowering plants are drastically reduced, breeders have strictly selected against early bolting. Consequently, genes controlling the phenological development have been mapped and cloned. First of all, the *B* locus controls early bolting. From this locus, the *BTC1* gene encoding a PSEUDO RESPONSE REGULATOR (PRR) has been cloned and functionally characterized [43]. In our study, the *BTC1* gene ranged from position 34,012,098 to 34,020,814 on chromosome 2. Accordingly, the strongest associations for bolting versus non-bolting were observed on chromosome 2. We found strong MTAs upstream of the *BTC1* gene but no MTAs within the gene. The recessive (biennial) *btc1* allele carries a 28 kb repeat-rich insertion in the promoter region, which is lacking from the dominant (annual) *BTC1* allele [43, 49]. We assume the marker polymorphisms are located within this insertion. However, Illumina short-reads used in our study do not allow genotyping of large insertions and deletions. Therefore, we propose long-range sequencing to localize the polymorphisms precisely. A yet unknown gene with a putative function in regulating circadian rhythm located within the associated region must be characterized in the future.

We identified 93 MTAs for Cercospora resistance spread across five chromosomes. Quantitative variation for Cercospora resistance was observed in the field, and the presence of MTAs on multiple chromosomes confirms the quantitative inheritance of CLS resistance. Within the strongest MTAs on chromosome 2 between position 30,827,268 and 31,330,290, we found a gene putatively encoding an actin-depolymerizing factor 4 (*ADF4*). Interestingly, a resistance QTL had been mapped on chromosome 2 in a previous study [22]. It is shown that ADFs play a crucial role in modulating the host’s cytoskeletal architecture and are involved in defense signaling during pathogen infection. The expression of *TaADF4*, a wheat homolog of *AtADF4*, has been demonstrated as necessary for resistance signaling in response to stripe rust (*Puccinia striiformis* f. sp. *Tritici*) and other biotic and abiotic stresses. Silencing *TaADF4* resulted in susceptibility to a virulent stripe rust race [50]. Interestingly, in *TaADF4*-silenced wheat plants, there was a reduction in jasmonic acid (JA) accumulation, leading the authors to propose that *TaADF4* plays a fundamental role in the signaling pathway, coordinating growth/defense signaling through JA and abscisic acid perception.

In Arabidopsis, *ADF4* is involved in the activation of defense signaling upon infection by the pathogen *Pseudomonas syringae* [51, 52]. Knockout mutants of *ADF4* exhibited reduced resistance. However, upon infection by the obligate biotroph powdery mildew fungus *Golovinomyces orontii*, the *ADF4* null mutant showed a significantly increased resistance. This enhanced resistance was linked to the accumulation of hydrogen peroxide and cell death specific to *G. orontii*-infected cells [53]. Interestingly, in oilseed rape (*Brassica napus*), pre-treatment with actin-depolymerizing drugs activated the SA pathway, leading to increased plant resistance to the hemibiotrophic fungal pathogen *Leptosphaeria maculans,* causing blackleg disease in *Brassica* crops [54].

We mapped a cytochrome P450 gene (*CYP94C1)* within a region on chromosome 7 associated with CLS infection. Among its many functions, cytochrome 450 controls JA oxidation. JA plays a crucial role apart from others in the response to abiotic and biotic stresses [55, 56]. It also promotes the synthesis of secondary metabolites such as flavonoids, glucosinolates, terpenoids, and phytoalexins [57, 58]. Thus, initiating plant resistance responses includes the biosynthesis of JA and the activation of a JA-dependent signaling cascade involving a group of transcription factors (TFs), among other events [59, 60]. Using a series of mutant and overexpression plant lines demonstrated that *CYP94B3*/*C1* are integral components of the fungus-induced jasmonate metabolic pathway. Moreover, *CYP94B3*/*C1* controls hormone oxidation status for signal attenuation and defines JA-Ile as a metabolic hub directing jasmonate profile complexity [61].

Within the associated region on chromosome 9, we found a cluster of genes putatively belonging to the family of NBS-LRR genes. Their function in plant-pathogen response has been explicitly documented as they can directly or indirectly detect pathogen proteins [62]. The NBS domain includes several kinds of nucleotide-binding kinases. Many plant pathogen resistance genes cloned so far encode NBS-LRR proteins, such as the tomato LRR-RLP gene *Cf-9,* which confers resistance to the biotrophic fungus *Cladosporium fulvum*, causing leaf mold [63]. Cf-9 recognizes the small cysteine-rich protein Avr9 secreted into the extracellular spaces of tomato leaves from *C. fulvum* and induces a wide range of defense responses [64, 65].

Both candidate genes *ADF4* and *CYP94C1* associated with *Cercospora* resistance deserve further attention to unveil the molecular background of CLS infections in *Beta* species. They are proposed to be major genes involved in resistance. Allele mining within our *Beta* diversity set and beyond is expected to detect new sequence variants. Breeding for CLS resistance is still challenging because the fungus has overcome most resistances introduced into varieties [66]. Therefore, pyramiding resistance alleles from the associated regions mapped in this study could help breed durable resistance.

## 5. Declaration of interests

The authors declare no competing interests.

## 6. Acknowledgments

We thank Brigitte Neidhardt-Olf, Bettina Rohardt, Ines Schütt, and Monika Bruisch for their technical assistance. We thank Prof. Anne-Katrin Mahlein, Annette Walter, and Celin Lachmann from IFZ, Göttingen, for supporting our *Cercospora* field trials. We gratefully acknowledge financial support from the German Science Foundation (Deutsche Forschungsgemeinschaft, DFG) – Project Number 400993799 (Project 2 within the Research Training Group 2501 Translational Evolutionary Research, https://gepris.dfg.de/gepris/projekt/400993799).

## Footnotes

1 https://www.oecd-ilibrary.org/agriculture-and-food/world-sugar-projections_e797df36-en

## 8. Supplementary data

### 8.1. Supplementary Tables

Supplementary Table 1: *Cercospora* disease index, ranging from 1 to 9, and description for each index value.

Supplementary Table 2: Phenotypic data from two field trials in 2021 and 2022 evaluated for different traits in this study. Seed codes and accession names marked as ‘Sample name’ used for whole genome sequencing are mentioned along with the phenotypic data from each accession.

Supplementary Table 3: Significant marker-trait associations for phenotypic traits evaluated in this study.

Supplementary Table 4: Summary statistics of significant marker-trait associations across the nine beet chromosomes. The ‘EL10’ genome was taken as a reference.

Supplementary Table 5: Genes 100-kb up- and downstream of SNPs significantly associated with hypocotyl GY (Green vs Yellow) color.

Supplementary Table 6: Genes 100-kb up- and downstream of SNPs significantly associated with hypocotyl GR (Green vs Red) color.

Supplementary Table 7: Genes 100-kb up- and downstream of SNPs significantly associated with root flesh color.

Supplementary Table 8: Genes 100-kb up- and downstream of SNPs significantly associated with external root color.

Supplementary Table 9: Genes 100-kb up- and downstream of SNPs significantly associated with leaf color.

Supplementary Table 10: Genes 100-kb up- and downstream of SNPs significantly associated with cambium ring color.

Supplementary Table 11: Genes 100-kb up- and downstream of SNPs significantly associated with petiole color.

Supplementary Table 12: Genes 100-kb up- and downstream of SNPs significantly associated with monogermity.

Supplementary Table 13: Genes 100-kb up- and downstream of SNPs significantly associated with bolting control.

Supplementary Table 14: Genes 100-kb up- and downstream of SNPs significantly associated with Cercospora resistance.

### 8.2. Supplementary Figures

Supplementary Figure 1: Photos showinge different phenotypic classes for traits investigated in the field. The photos were taken at the end of the field trial on September 29, 2021. The increasing level of CLS incidence on individual leaves in a highly susceptible sugar beet accession have been illustrated. Disease severity was evaluated using the Cercospora Disease Index, ranging from 1 to 9.

Supplementary Figure 2: *Cercospora* disease index evaluated at six different time points in 2021 (A) and 2022 (B). In each bar plot, the X-axis represents the individual accessions, sorted based on their *Cercospora* disease index, while the Y-axis shows the average *Cercospora* disease index value of each individual accession from three replication blocks, with the standard deviation indicated by the error bars. In total six ratings were perfomed in each year. Moving from left to right and top to bottom within the plots, the *Cercospora* disease ratings in 2021 were conducted at 90, 103, 118, 131, 146, and 159 days after sowing (DAS). In 2022, the ratings were conducted on 84, 98, 112, 125, 140, and 154 DAS.

Supplementary Figure 3: Manhattan plots for external root color, flesh color, and cambium ring color analyzed by genome-wide association study (GWAS). In each Manhattan plot, the X-axis represents the nine main ‘EL10’ chromosomes and unplaced scaffolds, which are labeled as 10, while the Y-axis illustrates the –log_10_(*p*) values. Two horizontal lines represent the Bonferroni thresholds of 7.02 (α=1) and 8.31 (α=0.05). The Manhattan plots were generated using the ‘qqman’ package in R. The *R* locus harbors the *CYP76AD1* gene, spanning from 49,286,874 to 49,291,710 bp, whereas the *Y* locus contains the *BvMYB1* gene, spanning from 51,654,941-51,658,812.

Supplementary Figure 4: Marker-trait associations for leaf color, and petiole color detected by a genome-wide association study. In each Manhattan plot, the X-axis represents the nine main ‘EL10’ chromosomes and unplaced scaffolds, which are labeled as 10, while the Y-axis illustrates the –log_10_(*p*) values. Two horizontal lines represent the Bonferroni thresholds of 7.02 (α=1) and 8.31 (α=0.05). The Manhattan plots were generated using the ‘qqman’ package in R. The *R* locus harbors the *CYP76AD1* gene, spanning the positions 49,286,874 to 49,291,710 bp, whereas the *Y* locus contains the *BvMYB1* gene, spans the positions 51,654,941 to 51,658,812.

Supplementary Figure 5: Genotypic and amino acid variation within the *WIP2* gene associated with monogermity in beets. (A) An IGV screenshot displaying the most significant SNP associated with monogermity. A red box highlights the position at 64,590,348 bp at the end of chromosome 4. The genotypes of monogerm, multigerm, and hybrid accessions are illustrated. (B) An alignment of cDNA sequences from the *WIP2* gene of multigerm beet, monogerm beet, and *Arabidopsis thaliana* is shown. Red vertical bars represent the percentage of conserved nucleotides among three *WIP2* cDNA sequences from multigerm beet, monogerm beet, and *Arabidopsis thaliana*. (C) A comparison of *WIP2* cDNA between monogerm and multigerm accessions, with the non-synonymous ‘missense’ mutation highlighted in the red box.

Supplementary Figure 6: Screenshots of InterPro and GO terms analysis showcase the BvWIP2 protein’s involvement in various biological, molecular, and cellular processes. The upper plot displays the complete BvWIP2 polypeptide, with predicted Classical Zinc finger C2H2 domains using InterPro scan. The lower plot is a zoomed-in version, highlighting the Zinc-finger domains between 319 and 442 amino acids.

Supplementary Figure 7: Q-Q plots of the *p*-values from GWAS analyses for monogermity, external root color, flesh color, cambium ring color, leaf color, and petiole color. In each Q-Q plot, the X-axis represents the expected –log_10_(*P*) values, while the Y-axis illustrates the observed –log_10_(*P*) values from the corresponding GWAS analysis. The red line indicates the deviation between expected and observed *p-*values. The Q-Q plots were created using the ‘qqman’ package in R.

Supplementary Figure 8: Q-Q plots of the *p*-values from GWAS analyses for hypocotyl color using green-yellow and green-red as extremes, bolting, and *Cercospora* resistance. In each Q-Q plot, the X-axis represents the expected –log_10_(*P*) values, while the Y-axis illustrates the observed –log_10_(*P*) values from the corresponding GWAS analysis. The red line indicates the deviation between expected and observed *p-*values. The Q-Q plots were created using the ‘qqman’ package in R.

## Notes

### Competing Interest Statement

The authors have declared no competing interest.

