## Supplementary Figures for "Uncovering Genes Essential in Domestication and Breeding of Sugar Beet"

### Slide 1
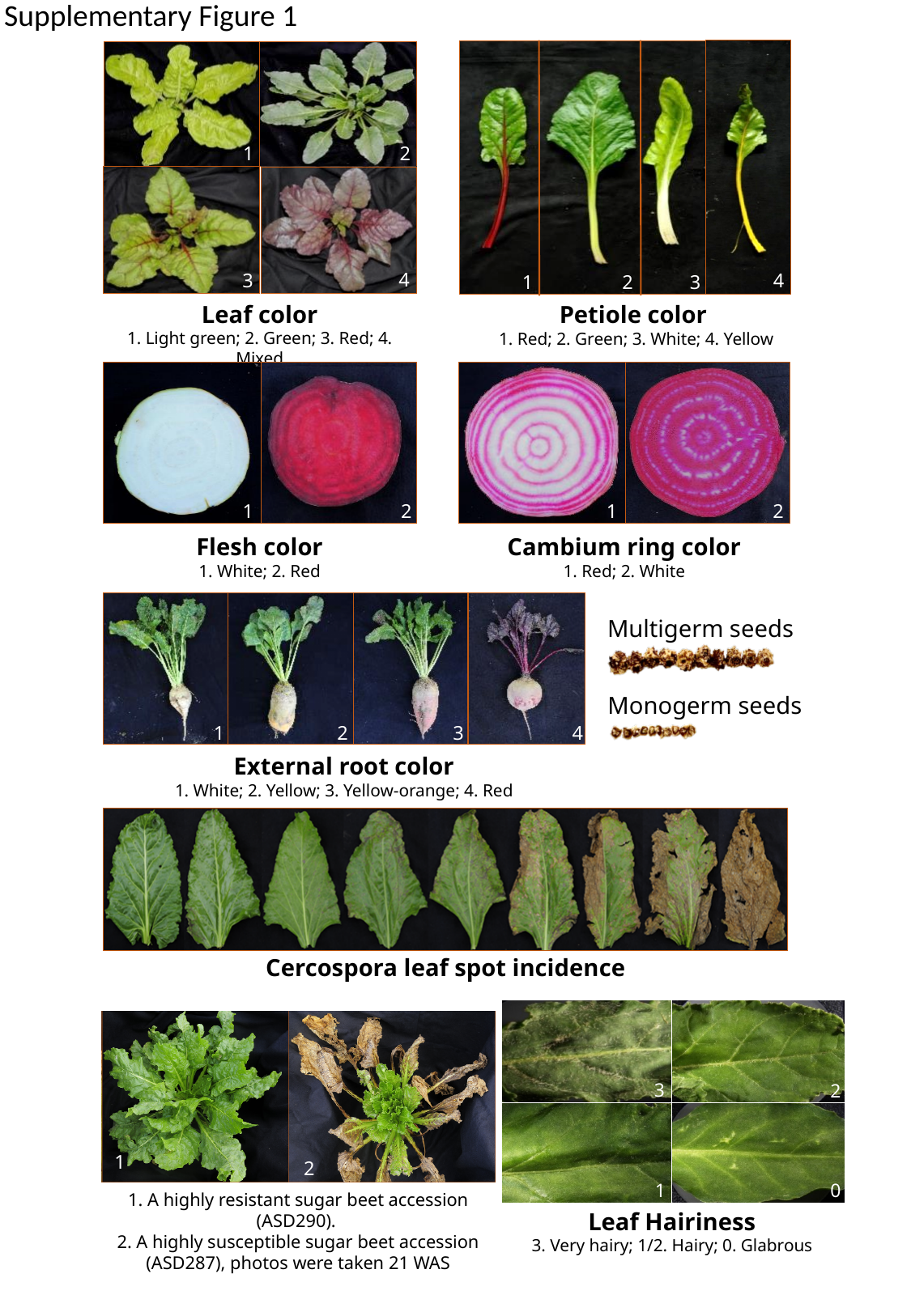

Supplementary Figure 1
1
2
3
Petiole color
1. Red; 2. Green; 3. White; 4. Yellow
4
1
2
4
3
Leaf color
1. Light green; 2. Green; 3. Red; 4. Mixed
Cambium ring color
1. Red; 2. White
1
2
Flesh color
1. White; 2. Red
1
2
1
2
3
4
External root color
1. White; 2. Yellow; 3. Yellow-orange; 4. Red
Multigerm seeds
Monogerm seeds
Cercospora leaf spot incidence
3
2
0
1
Leaf Hairiness
3. Very hairy; 1/2. Hairy; 0. Glabrous
1
2
1. A highly resistant sugar beet accession (ASD290).
2. A highly susceptible sugar beet accession (ASD287), photos were taken 21 WAS
1

### Slide 2
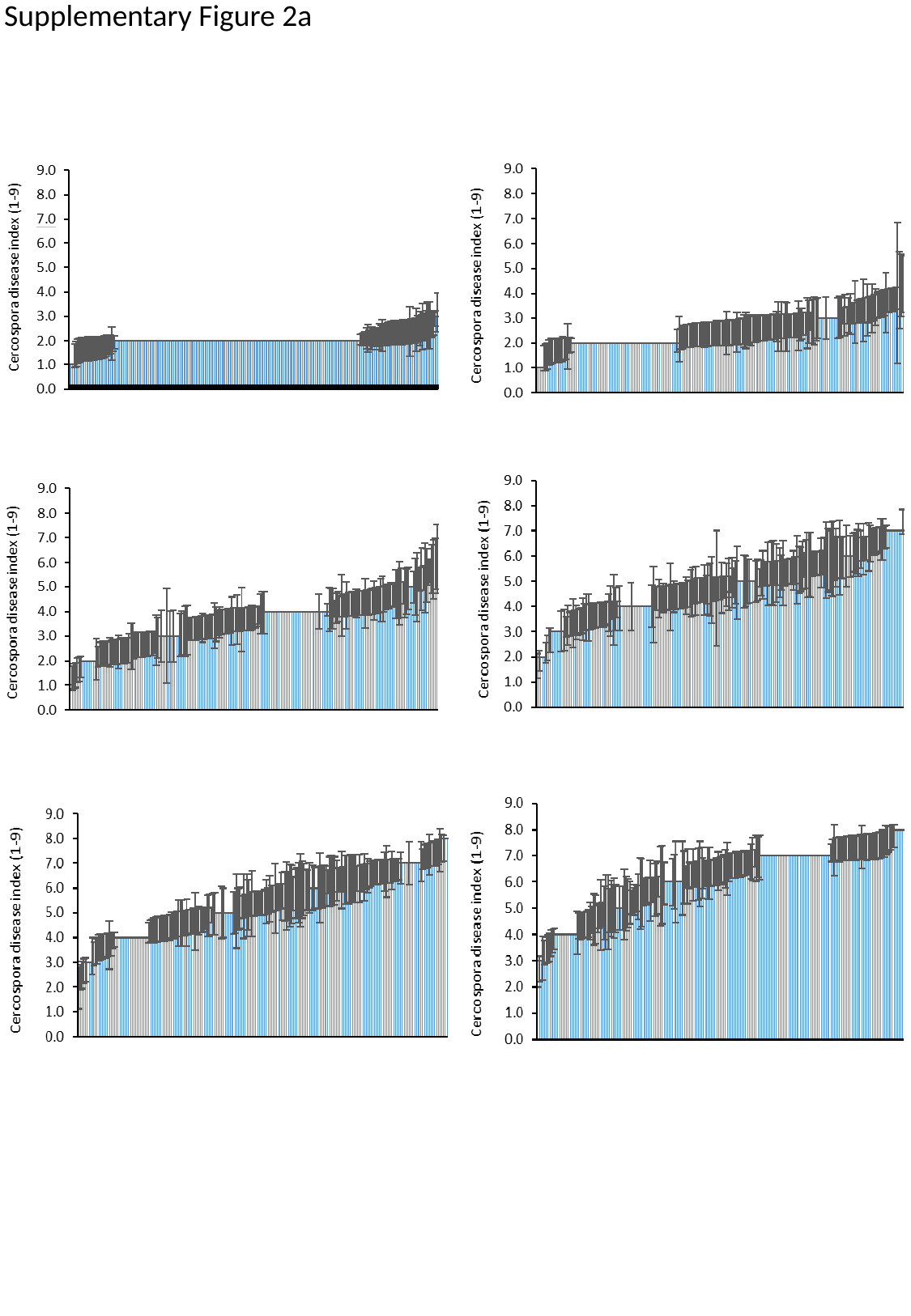

Supplementary Figure 2a

### Slide 3
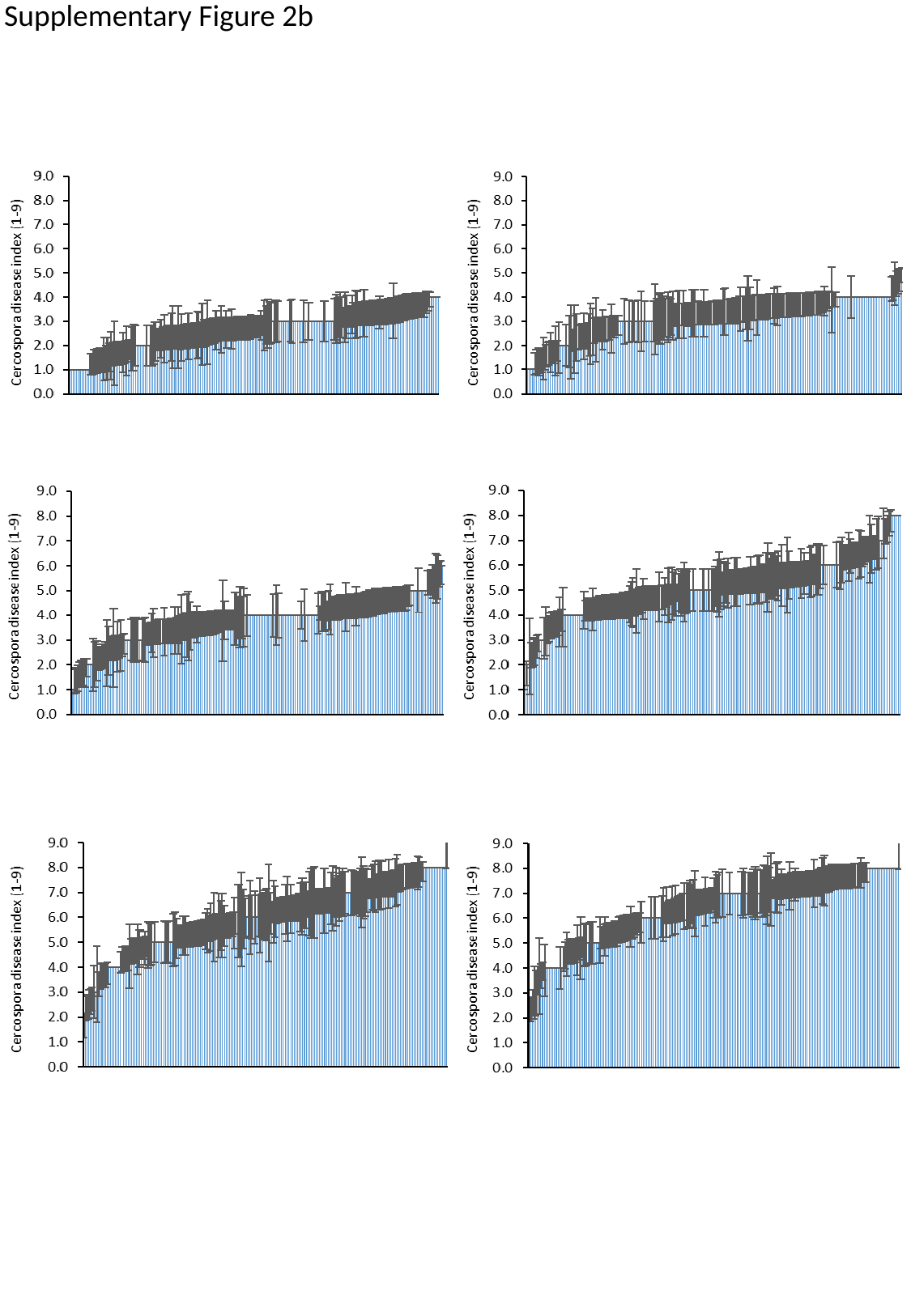

Supplementary Figure 2b

### Slide 4
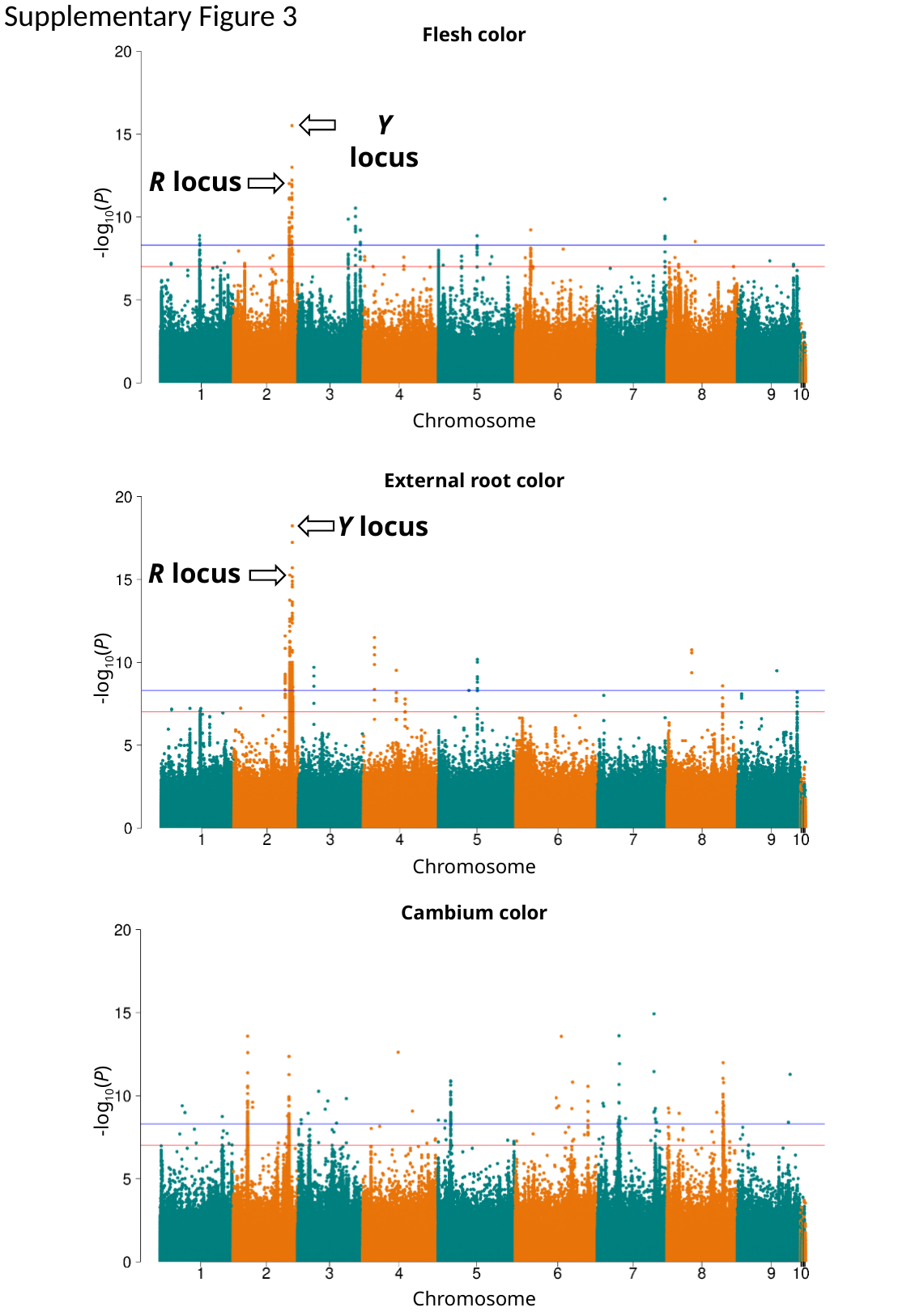

Supplementary Figure 3
Flesh color
-log_10_(P)
Chromosome
Y locus
R locus
black
External root color
Y locus
R locus
-log_10_(P)
Chromosome
Cambium color
-log_10_(P)
Chromosome

### Slide 5
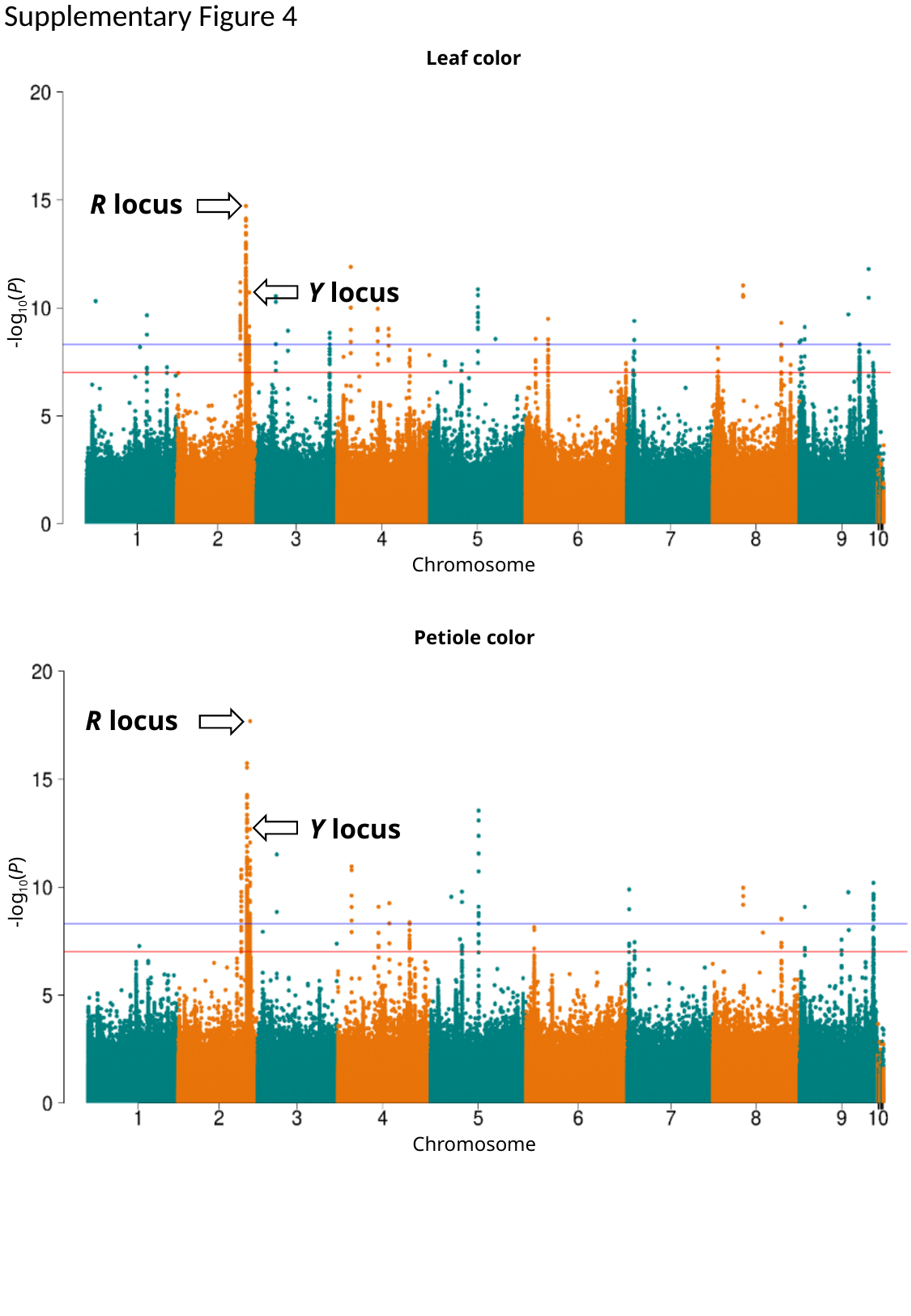

Supplementary Figure 4
Leaf color
-log_10_(P)
Chromosome
R locus
Y locus
Petiole color
-log_10_(P)
Chromosome
R locus
Y locus

### Slide 6
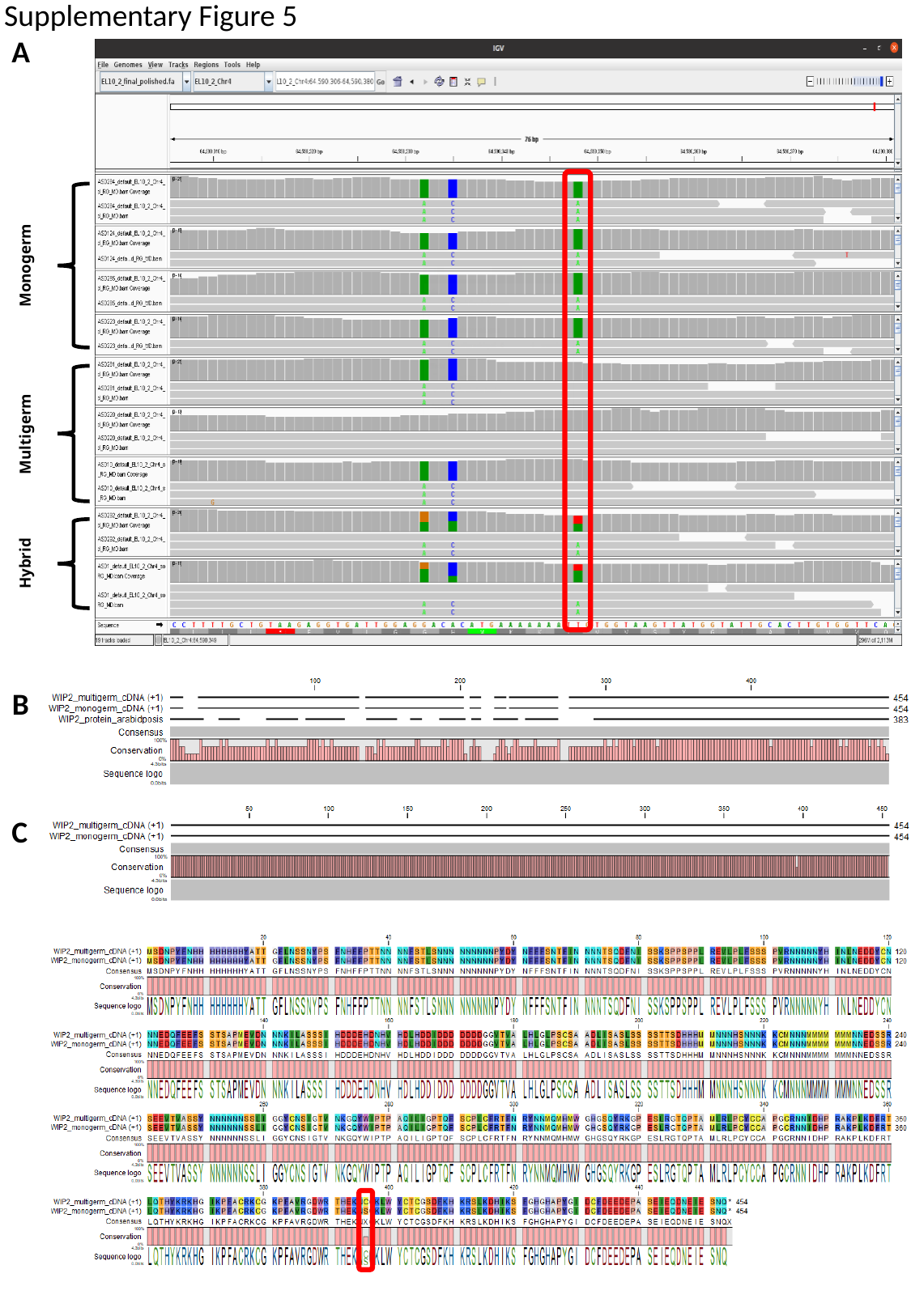

Supplementary Figure 5
A
Monogerm
Multigerm
Hybrid
B
C

### Slide 7
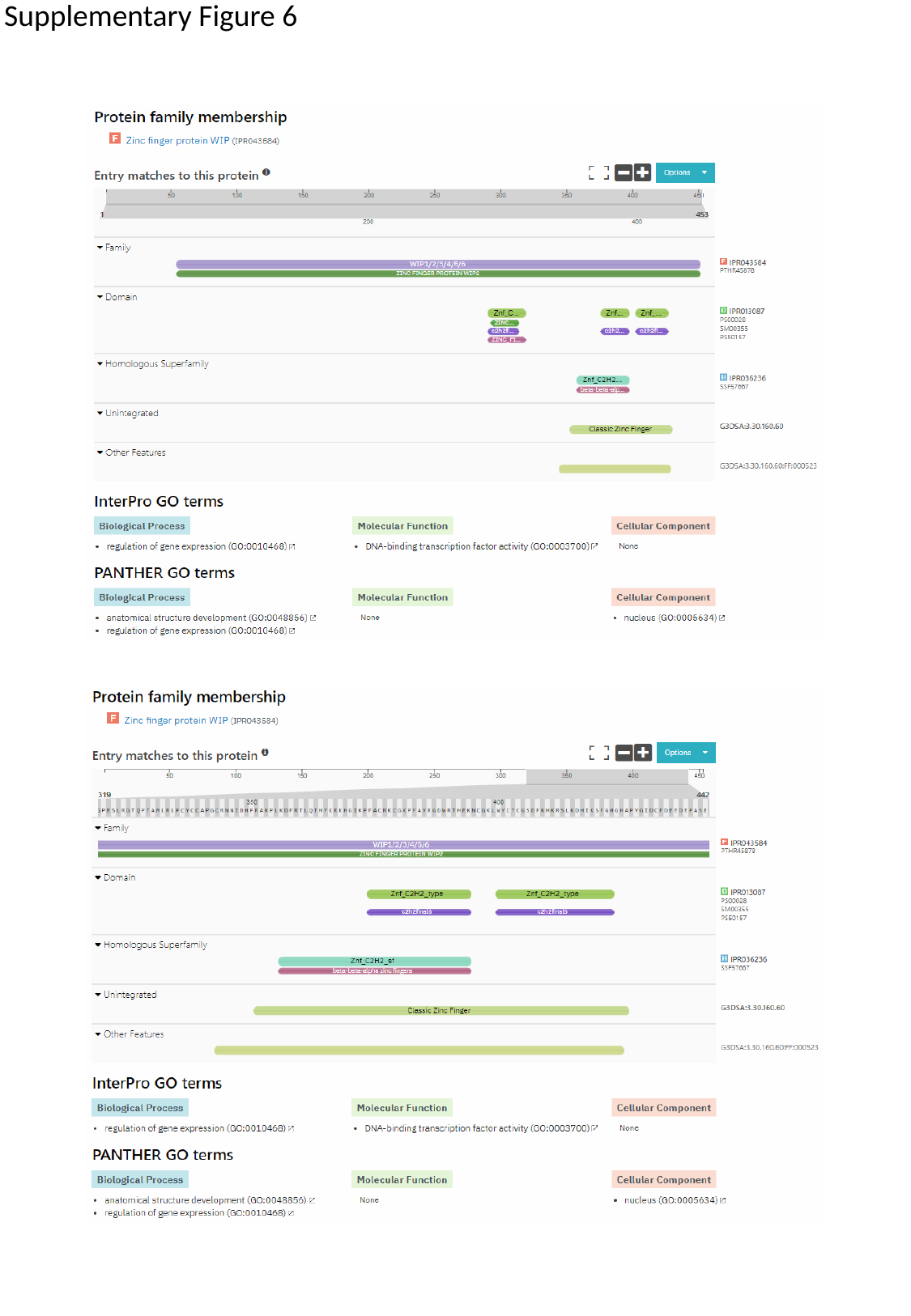

Supplementary Figure 6

### Slide 8
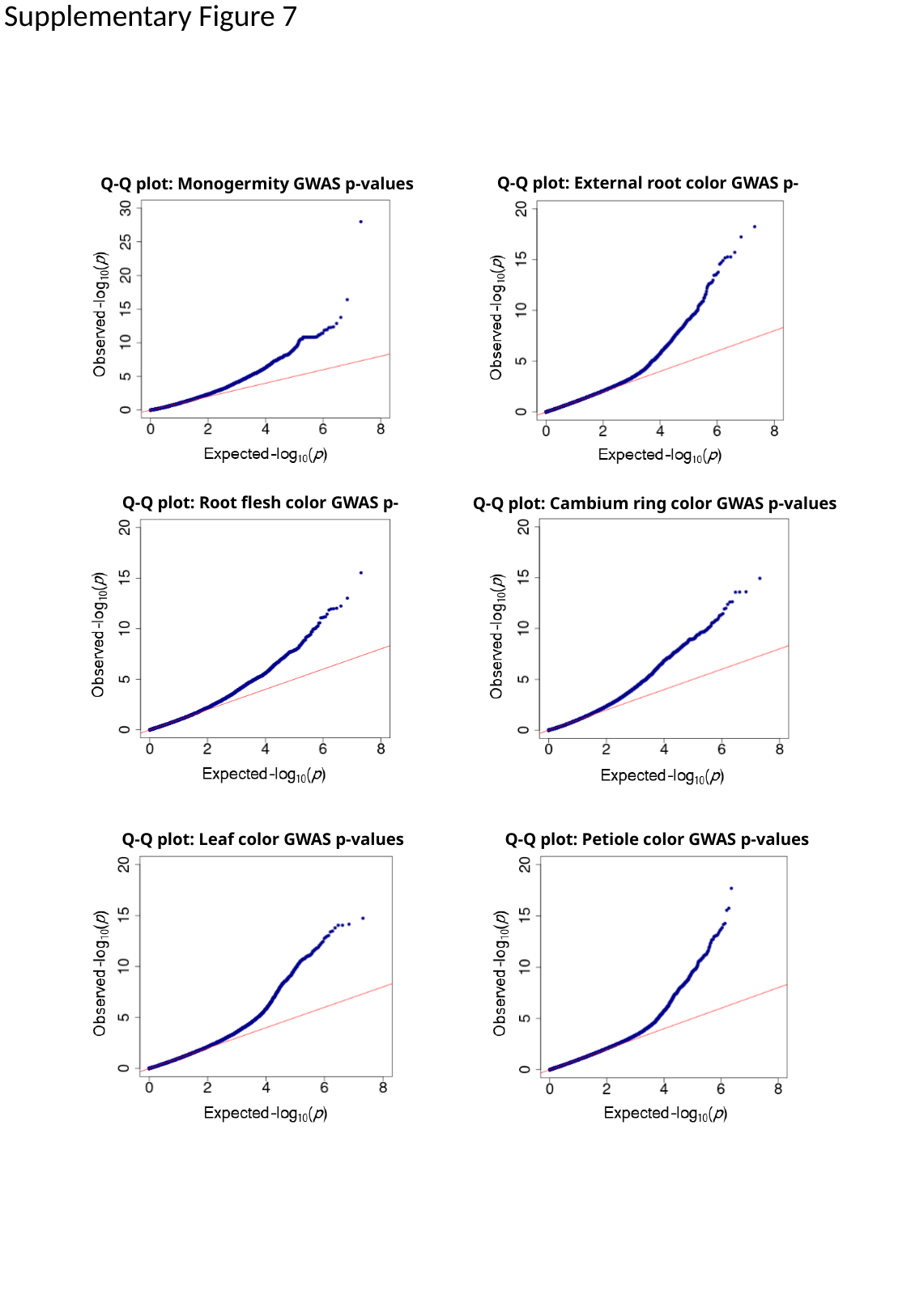

Supplementary Figure 7
Q-Q plot: External root color GWAS p-values
Q-Q plot: Monogermity GWAS p-values
Q-Q plot: Root flesh color GWAS p-values
Q-Q plot: Cambium ring color GWAS p-values
Q-Q plot: Leaf color GWAS p-values
Q-Q plot: Petiole color GWAS p-values

### Slide 9
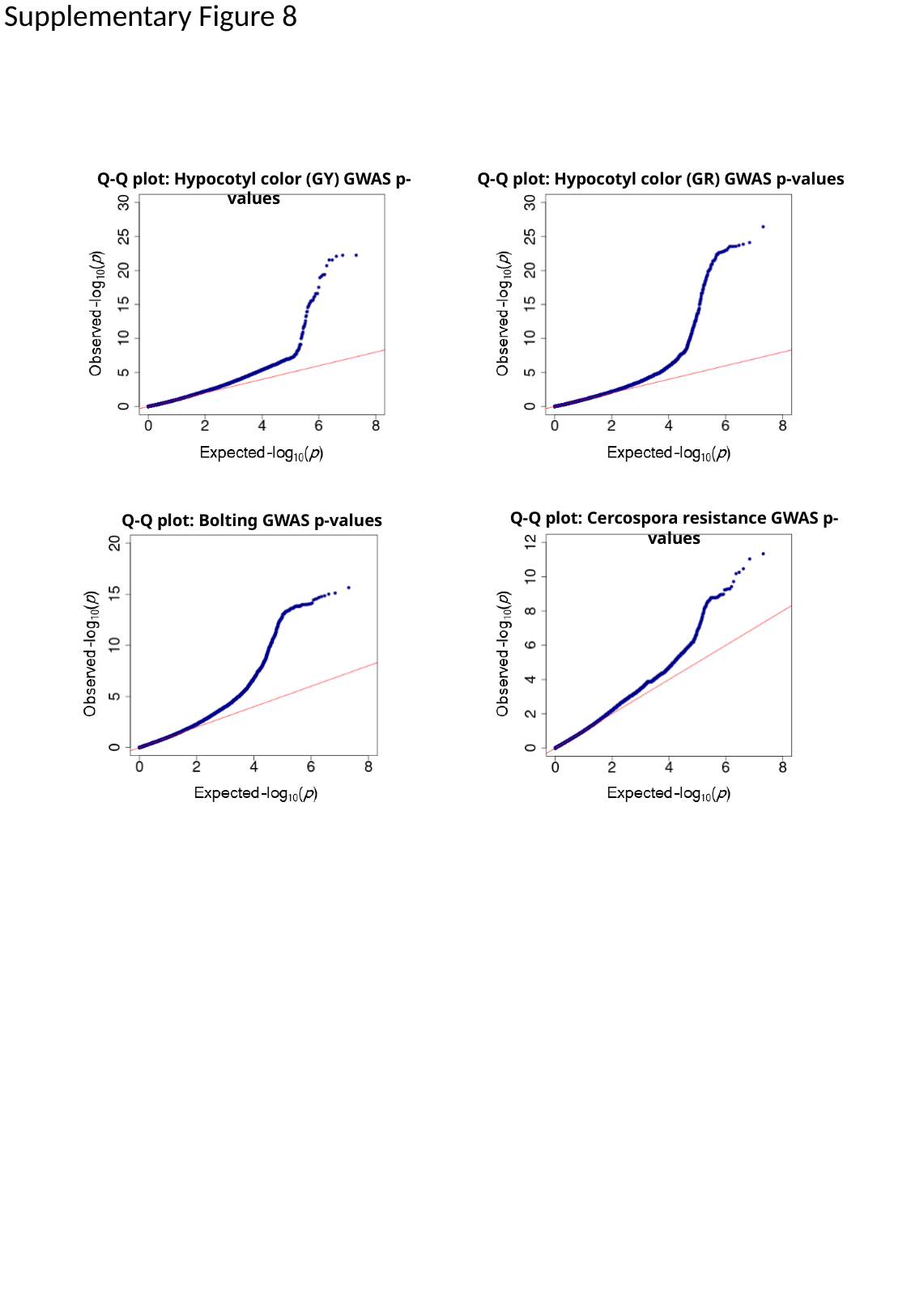

Supplementary Figure 8
Q-Q plot: Hypocotyl color (GY) GWAS p-values
Q-Q plot: Hypocotyl color (GR) GWAS p-values
Q-Q plot: Cercospora resistance GWAS p-values
Q-Q plot: Bolting GWAS p-values
